# Stress Coping Style Alters Functional Brain Network Activity to Acute Stressor

**DOI:** 10.64898/2025.12.05.692668

**Authors:** Jamie Corcoran, Kaylee Rushlau, Matthew R. Baker, Ryan Y. Wong

## Abstract

Consistent individual differences in behavior (e.g., personality types, stress coping styles) are a common occurrence across animal taxa. One hypothesis poses that the resulting constraints for within-individual behavioral variation may also lead to constraints for the evolution of behavior. With stress coping styles seen across taxa, it suggests a common underlying proximate mechanism. In this study we investigated neural activity patterns across the brain by quantifying immediate early gene expression in individuals with alternative stress coping styles in response to an acute stressor in zebrafish (*Danio rerio*). While immediate early gene expression levels of individual brain regions in the aversive brain network were similar across groups, functional network activity differed. There were several differences across groups including interactions between the basolateral amygdala and hippocampal homologs. One brain area that had many different connections across groups was the central gray. There were many differences involving central gray activity between proactive and reactive fish at baseline suggesting that baseline activity may prime for the reaction to stress. Collectively, baseline brain activity can predict behaviors, suggesting that these differences in brain interactions at baseline may be important for biasing behavioral responses to stressors that characterizes a stress coping style.

**Significance Statement:** It is well known that individuals cope with stress in different ways but the underlying mechanisms are not well understood. In this study we investigate how different brain regions interact under stress in animals of different stress coping styles. We investigated activity patterns across several brain regions in fish with a passive or active response to stress. We used an innovative statistical method that allowed us to compare how the brain regions function together under stress. The central gray was the most different between proactive and reactive groups, suggesting that this region is important for biasing the response to stressors. Further baseline functional connectivity differed the most between groups, suggesting that activity at baseline is important for biasing the response to stress mainly through connections with the central gray. Ultimately, functional neural network activity better explains stress behaviors than individual brain activity.

## Introduction

Behavioral and physiological variation between individuals in response to the same stressor can result in disparate but equally successful coping strategies. These differences are consistent within individuals, across contexts, and are potentially constrained by differences in brain activity (1, 2). One axis in which individuals can vary in response to stressors is their stress coping styles. Individuals with the proactive stress coping style tend to have a lower physiological stress response and respond actively to stress while individuals with a reactive stress coping style show the opposite (1). Identifying mechanisms underlying coping styles have focused on the neuroendocrine system (3, 4), select molecular pathways associated with the stress response (5, 6), and genome-wide expression patterns in the brain (7–10). However, it is less understood how larger brain networks process information that results in a biased behavioral and physiological response.

Variation in behavior can be attributed to differences in neural activity within a select few or a network of brain regions (11–14). Brain regions implicated in neuroendocrine stress responses across taxa (e.g., amygdala, limbic system, habenula, hippocampus, hypothalamus and other related regions) are key candidates for explaining variation in stress coping styles (15). While these brain regions can individually influence stress behavior, emergent properties of a functional neural network they belong to provides insight into how the information is processing and working towards a behavioral and physiological output (16). One network of interest is the aversive brain system in teleost fishes, which is composed of brain areas implicated in threat processing established by an increase in brain activity after exposure to alarm substance or other stressful stimuli (17). Important regions for stress and determining response to stress are the medial zone of the dorsal telencephalon (Dm), lateral zone of the dorsal telencephalon (Dl), ventral zone of the ventral telencephalon (Vv), superior zone of the ventral telencephalon (Vs), habenula (Hb), central gray (CG), preoptic area (POA), and caudal hypothalamus (Hc) (17).

Axonal projections to and from these regions as well as between these regions are involved in the emotional and fear encoding as well as regulation of the behavioral stress response (17–21). The complex interactions of these regions in the aversive brain system implies that the reaction to stress and threat is regulated by the interplay between multiple brain areas. Network activity has been implicated in individual differences in behavior (16). Therefore, differences in an established stress network may underlie the stress related behavioral differences between stress coping style.

In this study we quantified behavioral stress response and immediate early gene activity (*egr-1*) in 8 areas of the aversive brain system in proactive and reactive animals exposed to either an acute stressor or baseline control. We tested the hypothesis that differences in brain activity in the aversive brain system of the zebrafish are correlated with differences in stress behavior between personality type. We predicted that the interactions between different brain areas in this system, especially, the Dm, Dl, habenula, POA, caudal hypothalamus, and central gray, may explain the differences in stress coping style by controlling the level of perceived threat or reaction to the fear stimulus.

## Results

### Proactive fish displayed less stress behaviors compared to reactive fish

For both the proactive and reactive strains, we measured behavioral responses to an acute novelty stressor using a novel tank diving test. As expected, behavioral responses were significantly different between the strains (Figure 1). Proactive fish spent more time in the top half of the tank (t = 3.45, *p* = .001), spent less time frozen (t = -6.40, *p* = .000), traveled more distance (t = 7.36, *p* = .000), had more top transitions (t = 6.56, *p* = .000), less latency to enter the top half of the tank (t = -2.40, *p* = .012), and were faster (t = 7.33, *p* = .000) compared to reactive fish

**Figure 1.**
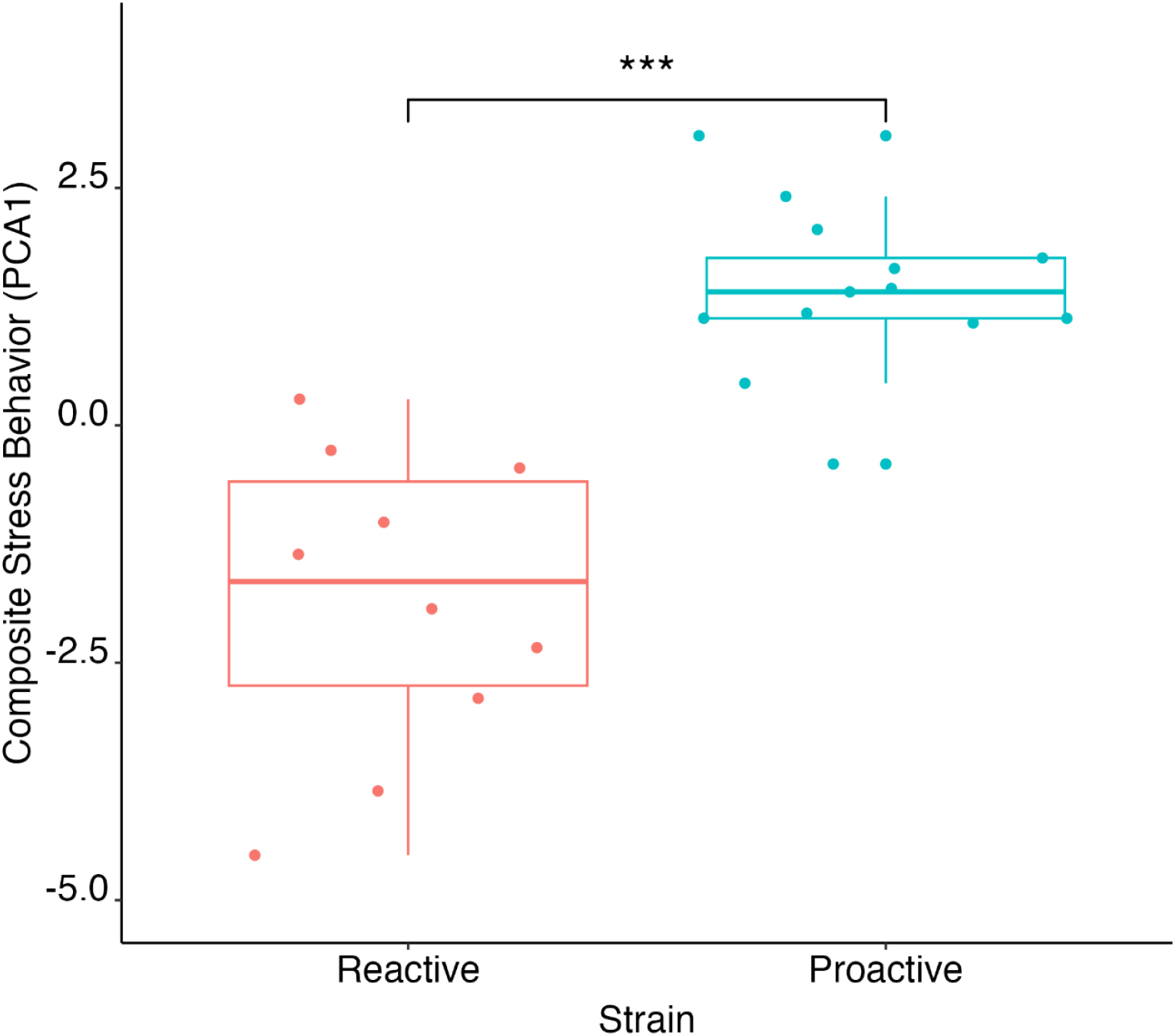
Comparison of PCA scores of behavior between proactive and reactive fish. There was a significant difference in scores between proactive and reactive fish (p<.001). Reactive fish are displayed in red and proactive fish in blue.

We also created a composite behavioral response by loading distance swam, velocity, top transitions, time spent in the top of the tank, latency to the top of the tank, and time spent moving, into a principal components analysis (PCA, Table S8). The first dimension accounted for 68.74% of the variance (eigenvalue = 4.13). All other dimensions had eigenvalues below one and were disregarded. For the first dimension velocity, distance, top transitions, time in top, and time spent moving loaded positively while the latency to the top zone loaded negatively. Using the composite score of each individual (PC score), we found that the proactive fish had significantly higher PC scores than reactive (t = -5.87, p = .0005).

### Creation of structural equation models (SEM) for stressed and baseline groups

We used SEM to explore differences in how *egr-1* expression across the eight measured brain regions relate to each other between stressed and baseline and proactive and reactive fish. First the data was partitioned into stressed and baseline groups where we assessed the optimal models for each group. As many of the communications in the brain that are thought to promote the response to stress in the aversive brain system are from forebrain regions to hindbrain regions (17), we then adjusted the model by starting with forebrain to hindbrain regions and testing other associations by seeing if there was a significant change in chi squared values (Figure 2). The optimal model for the stressed group was assessed first (χ2(25) = 14.04, p = .17, CFI = 0.96, RMSEA = .17, SRMR = .05) and then the same model was applied to the control group (χ2(31) = 7.77, p = .17, CFI = 0.94, RMSEA = .11, SRMR = .06). To test within coping style between baseline and stressed we used the same model. For the proactive group the initial fit was (χ2(32) = 8.57, p = .13, CFI = 0.93, RMSEA = .13, SRMR = .07). For the reactive group the initial fit was not acceptable (χ2(25) =14.62, p = .15, CFI =0.92, RMSEA = .20, SRMR = .05) so we tested another model. The model we tested had good fit ((χ2(25) = 17.37, p = .30, CFI = 0.97, RMSEA = .11, SRMR = .07).

**Figure 2.**
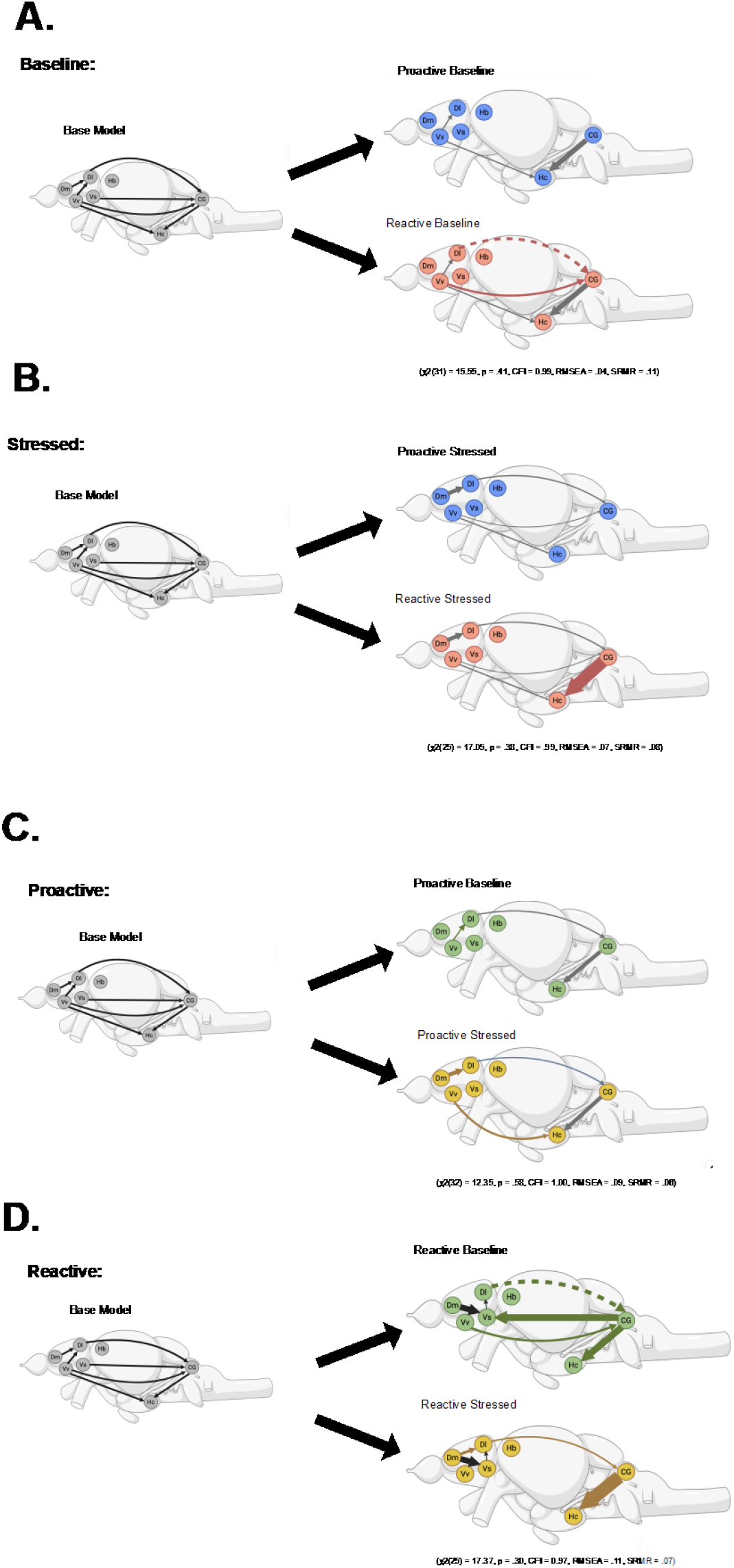
SEM models of stressed and baseline fish compared between proactive and reactive fish and proactive and reactive fish compared across stressed and baseline. (A) depicts the comparison between proactive and reactive baseline fish with proactive in blue and reactive in red. (B) depicts the comparison between proactive and reactive stressed fish with proactive in blue and reactive in red. (C) depicts the comparison between proactive baseline and stressed fish with baseline in green and stressed in yellow. (D) depicts the comparison between reactive baseline and stressed with baseline in green and stressed in yellow. The base model is depicted with all brain regions in gray with the fit statistics underneath. The colored lines indicate differences between the two models tested. The gray lines indicate that the interaction is the same between proactive and reactive fish The dotted lines indicate a negative relationship and solid lines indicate a positive relationship. The weight of the lines indicate the degree of interaction, defined by the B value. The image was created using BioRender.

### Differences in interactions of egr-1 expression in brain areas across stress coping styles and treatment

Within each group we split the models by stress coping styles and constrained the paths to test differences in interactions across strain (Figure 2A, 2B). In the stressed group the interaction that was significantly different between coping styles was the effect of *egr-1* levels in CG on Hc (only in reactive) (Table 1). The fit statistics were acceptable (χ2(25) = 17.05, p = .38, CFI = .99, RMSEA = .07, SRMR = .08). Comparing between strains in the baseline group the interactions that were significantly different between groups were the effect of the Dl on the CG (only in reactive) and the effect of the Vv on the CG (only in reactive). Fit statistics were acceptable (χ2(31) = 15.55, p = .41, CFI = 0.99, RMSEA = .04, SRMR = .11).

We also tested within coping style across baseline and stressed conditions to investigate how the *egr-1* expression networks change when stressed within each coping style (Figure 2C, 2D). The fit for the proactive group when split by group and constrained was (χ2(32) = 12.35, p = .58, CFI = 1.00, RMSEA = .09, SRMR = .06). In the proactive group the interactions that were different between baseline and stressed conditions were the effect of *egr-1* expression levels in the Vv on Dl (only in baseline condition), Dm on the Dl (only in stressed condition) and Vv on Hc (only in stressed condition). The fit for the reactive group after being split and constrained was (χ2(28) = 17.37, p = .30, CFI = 0.97, RMSEA = .11, SRMR = .07). For the reactive group the interactions that were different across baseline and stressed conditions were the effect of the Dm on the Dl (only in stressed), Vs on Dl (only in baseline), CG on Vs (only in baseline), CG on Hc (stronger in stressed), Vv on Hc (only in baseline), Dl on CG (negative in baseline, positive in stressed), Vv on CG (only in baseline).

While there were differences in the patterns of network activity across groups, 2-way ANOVAs of *egr-1* expression of individual brain regions did not show any differences across strain or treatment. To investigate whether changes in the magnitude of behavior relates to *egr-1* expression in each brain area and how neural activity patterns correlate between brain regions, we conducted Spearman’s correlations with Benjamani-Hochberg correction. There were no significant correlations between brain areas and behavior. There were a few correlations between brain areas with a very strong correlation between the Dm and Dl (*r* = .87, *p* < .05) (Supplementary Table 5).

## Discussion

In this study we investigated how differences in neural activity patterns may bias behavioral responses to a stressor between proactive and reactive fish. Individual behaviors and PCA scores showed that, as expected, the proactive fish had lower stress-related behaviors than reactive fish. While there were no differences in *egr-1* expression across groups in individual brain areas, we identified a difference between treatment and stress coping styles when examining functional neural activity networks. These differences point to key network interaction shifts that may explain biases in behavior between the coping styles.

There were no significant differences in neural activity in any brain region between any of the groups. These results suggest that neural activity of a single brain region cannot explain behavioral differences between the stress coping styles in response to a stressor. However, examining functional neural activity patterns across the 8 brain regions showed differences in network activity between groups (Tables S1-4). In reactive stressed fish there was a strong connection from the Dm to Dl that is not present in baseline fish (Figure 2D). This connection is likely due to Dm and Dl’s (homologs to the mammalian basolateral amygdala and hippocampus, respectively) involvement in processing and encoding internal and external stressor information (22–24). While the connection between the CG and Hc was present in both baseline and stressed conditions, the connection was stronger (higher B value) in stressed conditions. This increased connection strength is likely due to the central gray regulating the neuroendocrine stress axis (25). Interestingly, the reactive baseline fish had the most connections in the aversive brain system, especially involving the central gray (Figure 2D). These were positive connections from the CG to the Vs, and Vv to the CG. While a prior study found that the inhibitory connection between the medial prefrontal cortex and the central gray decreased passive coping style (freezing behavior) in mice (26), the connections at baseline in our reactive fish are mainly positive. Similarly, connections between regions like the Vs and habenula to the central gray are thought to mediate the response to threat (17, 27). Our study supports this idea but also suggests that the activity at baseline is biasing certain responses to stress. We also have a connection from the Dl to the CG in stressed fish. Interestingly the negative connection is only seen in reactive baseline fish while the positive connection is present in every other group. It is unclear what function this negative connection is performing but its presence adds evidence to the unique network activity in reactive baseline fish. This unique activity in the aversive brain system in reactive fish at baseline may explain the differences in stress behavior between proactive and reactive individuals.

There were differences in network activity between baseline and stressed proactive fish (Figure 2C). The connection between the Vv and the hypothalamus was only observed in proactive stressed fish. The Vv shows similarities to the lateral septal complex in mammals (28). In mammals, the lateral septum has been seen to be involved in stress response. Altering lateral septum activity altered stress coping, where increasing activity increased active coping and decreased HPA axis and decreasing activity had the opposite effect (29). In teleosts, the Vv may be performing a similar function where projections to the hypothalamus are increasing active coping. In proactive fish at baseline there is no interaction between the Dm and Dl while there is a positive interaction under stressed conditions. The positive interaction under stress may be due to both regions increasing serotonergic activity in response to stressful stimuli (22). Stress has been seen to increase activity between the amygdala and hippocampus (30). There is also a small connection between the Vv and Dl in baseline proactive fish that is not present in stressed proactive fish. Connectivity between the Vv and Dl is normal and does not appear to be related to stress (31). In the proactive fish there is less connectivity in the aversive brain system especially at baseline.

There was only one difference between reactive and proactive stressed fish (Figure 2B). In reactive stressed fish there is a connection from the CG to the Hc that is not in proactive stressed fish. This connection is likely activating the HPA axis in reactive fish. We hypothesize that this connection is the central gray regulating the neuroendocrine stress axis (25). It’s presence only in reactive fish could explain their faster cortisol release rate in response to a novelty stressor (4). The central gray may be an important region for constraining response to threat at baseline and for determining the active response to stress under threat. These differences in connectivity seem to bias towards passive or active coping style and explain these individual differences in response to stress.

At baseline we observed varying directions of interactions between CG and the forebrain in the reactive baseline group and none in the proactive baseline group (Figure 2A). While we did not examine specific subregions of the central gray, another study showed that different subpopulations of neurons in the central gray mediate different responses to stress through interactions with other regions of the brain (25). The connections at baseline in reactive fish are likely mediating responses to stress through interactions with many different brain regions. The central gray is important for determining the response to stress (25). Multiple studies have shown that altering activity in the central gray alters stress response (26, 32). A more specific study into the parts of the CG and how they interact with other regions to affect coping style could clarify these results

Intriguingly, the largest difference between coping style was in baseline activity. Other studies also see differences in resting state activity between individuals with different behavioral traits (33–37), suggesting that resting state activity plays an important role in determining behavior possibly due to spontaneous brain activity that is modelling and predicting outcomes (38). In the present study we posit that spontaneous activity would differ between stress coping styles where reactive individuals might predict more threat and that a freezing response would provide a better outcome where proactive individuals would predict the opposite. Each stress coping style could be acting with different models of the environment, shown by their differences in baseline brain activity, that lead to their differences in behavioral stress responses.

## Materials and Methods

### Subjects

We used zebrafish from lines selectively bred to display the proactive (n = 30) and reactive (n = 30) stress coping styles (39). Across six different stress and anxiety-like behavioral assays and across time, the reactive line exhibits higher stress and anxiety-related behaviors compared to the proactive line (39, 40). The reactive fish also shows faster release of cortisol under stress and distinct brain neurotranscriptome profile at baseline compared to the proactive line (4, 8). In this study fish were 12 generations removed from wild caught individuals located approximately 60 km from Kolkata, India. We kept fish together in mixed-sex 40 liter tanks on a recirculating water system with solid filtration on a 14:10 L/D cycle at a temperature of 27 °C. We fed fish twice a day with Tetramin Tropical Flakes (Tetra, USA).

### Procedures Stress Assay

The novel tank diving test (NTDT) is a standard behavioral assay to measure stress from a variety of stressors (41–45). Following established protocols, we video-recorded individual fish in a 30-minute NTDT (4, 39, 46, 47). We quantified time spent in the top and bottom half of the tank measured along with number of transitions, distance traveled, velocity, and time spent frozen using commercial software (Ethovision version 17.5.1718). See Supplementary Table 8 for additional details.

### In situ Hybridization

We collected and stored tissue samples at -80°C immediately after the NTDT for stressed groups and directly after catching from home tanks for baseline groups. Tissue samples were sectioned in four serial series at 16 µm on a cryostat (Thermoscientific, HM525 NX). Tissue fixation parameters, probe synthesis, ISH conditions, and quantification were based on established protocols(48, 49). In brief, we used digoxigenin (DIG)-labelled probe for the *egr-1* gene.

Riboprobes showed specific binding with high expression using the antisense probe and no expression in the sense condition. We identified the medial zone of the dorsal telencephalon, the lateral zone of the dorsal telencephalon, habenula, preoptic area, supracommisural zone of the ventral telencephalon, ventral zone of the ventral telencephalon, caudal hypothalamus, and central gray using the zebrafish brain atlas (50). Across individuals within each brain region we quantified the mean optical density (OD) of *egr-1* expression (Nikon NIS Elements Version 4.6) from images captured with a DS-Qi2 monochrome camera (Nikon). For each slide, we normalized the mean intensity of all measures to the background (mean intensity of slide area not containing tissue), which produced a fractional transmittance value for each brain region in each section.

Fractional transmittance was mathematically converted to optical density by the equation OD = 2 – log(fractional transmittance). The researcher was blind to the strain and treatment data when analyzing the brain regions. See Supplementary Table 6 for additional details.

### Statistical Analyses

We used a t-test to compare the behaviors in the NTDT between proactive and reactive individuals. We also performed a PCA on the behavior data. Then we conducted a t-test on the eigenvalues comparing across stress coping style. We performed a two-way ANOVA with stress and treatment to look at any differences in *egr-1* expression in each of the brain areas. ANOVA and t-test analyses were performed in R.

We used structural equation modelling to investigate the differences in brain area interactions between proactive and reactive individuals under stressed and not stressed conditions. We used a maximum likelihood estimation with robust standard errors due to not having normally distributed data. First we split the data by baseline and stressed to allow for comparisons of proactive and reactive individuals within these groups. We tested the stressed group first and created an optimal model. Then we grouped the model by strain to compare differences across coping style in the stressed group. We constrained, made equal, each path individually and if the change in chi squared was less than 3.84 we kept the paths equal across the group being compared. The same model was used for all other comparisons except for the reactive group. We used the same procedure to compare within the proactive group across control and treatment and within the reactive group between control and treatment. All of the models were the same except for the model for the reactive fish. We excluded fish with multiple brain areas with missing values. We excluded the preoptic area (POA) from all models due to a large amount of missing values. We tested models using R lavaan package (51).

## Acknowledgments

We thank Daniel Revers, Elizabeth Stone, Amber Park, and Nagat Mohammad for help with fish husbandry. We are grateful to Alexander Goodman, Rebecca Patterson, Sean Bresnahan, and Kylie Cullen for helpful discussions and technical assistance. This project was funded by the National Institutes of Health (R15MH113074), Nebraska EPSCoR First Award (OIA-1557417), Nebraska Research Initiative, and UNO Office of Creative and Research Activities, and startup funds to RYW.

## Notes

### Competing Interest Statement

The authors have declared no competing interest.

