## Supplemental Tables, Figures, and Methods for "Stress Coping Style Alters Functional Brain Network Activity to Acute Stressor"


Jamie Corcoran[1]

Kaylee Rushlau[2]

Matthew R. Baker[2,3]

Ryan Y. Wong[1,2]

1. Department of Psychology, University of Nebraska at Omaha, Omaha, NE
2. Department of Biology, University of Nebraska at Omaha, Omaha, NE
3. Department of Neurosurgery, Mayo Clinic, Rochester, MN

*Ryan Y. Wong

**Supplementary Table 1. SEM Results for each model comparison of Baseline Proactive vs reactive**

**A. Proactive**

| Hub | Brain Area | Constrained | B | SE | *p* | β | $R^2$ |
|---|---|---|---|---|---|---|---|
| Dl | | | | | | | 0.27 |
| | Dm | Yes | -0.23 | 0.18 | 0.20 | -0.23 | |
| | Vv | Yes | 0.28 | 0.09 | 0.002 | 0.28 | |
| Hc | | | | | | | |
| | CG | Yes | 1.21 | 0.24 | <0.01 | 1.21 | 0.4 |
| | Vv | Yes | 0.23 | 0.11 | 0.04 | 0.23 | |
| CG | | | | | | | |
| | Dl | No | 0.22 | 0.15 | 0.14 | 0.22 | 0.08 |
| | Vv | No | -0.09 | 0.11 | 0.42 | -0.09 | |
| | Vs | Yes | 0.02 | 0.04 | 0.68 | 0.02 | |

*Note. N* = 16

**B. Reactive**

| Hub | Brain Area | Constrained | B | SE | *p* | β | $R^2$ |
|---|---|---|---|---|---|---|---|
| Dl | | | | | | | 0.29 |
| | Dm | Yes | -0.23 | 0.18 | 0.20 | -0.23 | |
| | Vv | Yes | 0.28 | 0.09 | 0.002 | 0.28 | |
| Hc | | | | | | | |
| | CG | Yes | 1.21 | 0.24 | <0.01 | 1.21 | 0.62 |
| | Vv | Yes | 0.23 | 0.11 | 0.04 | 0.23 | |
| CG | | | | | | | |
| | Dl | No | -0.67 | 0.29 | 0.02 | -0.67 | 0.48 |
| | Vv | No | 0.48 | 0.19 | 0.01 | 0.48 | |
| | Vs | Yes | 0.02 | 0.04 | 0.68 | 0.02 | |

*Note. N* = 15

**Supplementary Table 2. SEM Results for each model comparison of Stressed Proactive vs reactive**

**A. Proactive**

| Hub | Brain Area | Constrained | B | SE | *p* | β | $R^2$ |
|---|---|---|---|---|---|---|---|
| Dl | | | | | | | 0.88 |
| | Dm | Yes | 0.91 | 0.09 | <.01 | 0.95 | |
| | Vv | Yes | -0.02 | 0.04 | 0.69 | -0.02 | |
| Hc | | | | | | | 0.37 |
| | CG | No | 0.54 | 0.49 | 0.27 | 0.26 | |
| | Vv | Yes | 0.29 | 0.08 | <.01 | 0.54 | |
| CG | | | | | | | 0.53 |
| | Dl | Yes | 0.33 | 0.05 | <.01 | 0.87 | |
| | Vv | Yes | -0.06 | 0.02 | <.01 | -0.23 | |
| | Vs | Yes | -0.04 | 0.05 | 0.39 | -0.16 | |

*Note. N* = 15

**B. Reactive**

| Hub | Brain Area | Constrained | B | SE | *p* | β |
|---|---|---|---|---|---|---|
| Dl | | | | | | |
| | Dm | Yes | 0.91 | 0.09 | <.01 | 0.93 |
| | Vv | Yes | -0.02 | 0.04 | 0.69 | -0.04 |
| Hc | | | | | | |
| | CG | No | 3.91 | 0.45 | <.01 | 0.88 |
| | Vv | Yes | 0.29 | 0.08 | <.01 | 0.31 |
| CG | | | | | | |
| | Dl | Yes | 0.33 | 0.05 | <.01 | 0.62 |
| | Vv | Yes | -0.06 | 0.02 | <.01 | -0.29 |
| | Vs | Yes | -0.04 | 0.05 | 0.39 | -0.11 |

*Note. N* = 10

**Supplementary Table 3. SEM Results for each model comparison of Proactive Baseline vs stressed**

| Hub | Brain Area | Constrained | B | SE | *p* | β | $R^2$ |
|---|---|---|---|---|---|---|---|
| Dl | | | | | | | 0.43 |
| | Dm | No | -0.47 | 0.27 | 0.09 | -0.36 | |
| | Vv | No | 0.31 | 0.09 | <.01 | 0.43 | |
| Hc | | | | | | | 0.29 |
| | CG | Yes | 0.94 | 0.31 | <.01 | 0.54 | |
| | Vv | No | 0.05 | 0.16 | 0.77 | 0.06 | |
| CG | | | | | | | 0.17 |
| | Dl | Yes | 0.29 | 0.05 | <.01 | 0.44 | |
| | Vv | Yes | -0.05 | 0.02 | 0.07 | -0.09 | |
| | Vs | Yes | -0.04 | 0.03 | 0.14 | -0.22 | |

**A. Baseline**

*Note. N* = 17

**B. Stressed**

| Hub | Brain Area | Constrained | B | SE | *p* | β | $R^2$ |
|---|---|---|---|---|---|---|---|
| Dl | | | | | | | 0.89 |
| | Dm | No | 0.99 | 0.12 | <.01 | 0.96 | |
| | Vv | No | -0.01 | 0.06 | 0.84 | -0.02 | |
| Hc | | | | | | | 0.63 |
| | CG | Yes | 0.94 | 0.31 | <.01 | 0.54 | |
| | Vv | No | 0.50 | 0.16 | <.01 | 0.50 | |
| CG | | | | | | | 0.49 |
| | Dl | Yes | 0.29 | 0.05 | <.01 | 0.44 | |
| | Vv | Yes | -0.05 | 0.02 | 0.07 | -0.09 | |
| | Vs | Yes | -0.04 | 0.03 | 0.14 | -0.22 | |

*Note. N* = 15

**Supplementary Table 4. SEM Results for Reactive Baseline vs Stressed**

**A. Baseline**

| Hub | Brain Area | Constrained | B | SE | *p* | β | $R^2$ |
|---|---|---|---|---|---|---|---|
| Dl | | | | | | | 0.02 |
| | Dm | No | -0.04 | 0.22 | 0.84 | -0.05 | |
| | Vs | Yes | 0.18 | 0.08 | 0.03 | 0.80 | |
| Vs | | | | | | | 0.07 |
| | CG | No | 2.34 | 0.90 | 0.01 | 0.75 | |
| | Dm | Yes | 1.12 | 0.15 | <.01 | 0.26 | |
| Hc | | | | | | | 0.69 |
| | CG | No | 1.38 | 0.25 | <.01 | 0.73 | |
| | Vv | Yes | 0.20 | 0.16 | 0.20 | 0.16 | |
| CG | | | | | | | 0.47 |
| | Dl | No | -1.05 | 0.33 | <.01 | -0.73 | |
| | Vv | No | 0.57 | 0.18 | <.01 | 0.85 | |

*Note. N* = 15

**B. Stressed**

| Hub | Brain Area | Constrained | B | SE | *p* | β | $R^2$ |
|---|---|---|---|---|---|---|---|
| Dl | | | | | | | 0.84 |
| | Dm | No | 0.59 | 0.12 | <.01 | 0.69 | |
| | Vs | Yes | 0.18 | 0.08 | 0.03 | 0.80 | |
| Vs | | | | | | | |
| | CG | No | -0.16 | 0.47 | 0.74 | -0.06 | 0.72 |
| | Dm | Yes | 1.12 | 0.15 | <.01 | 0.88 | |
| Hc | | | | | | | 0.71 |
| | CG | No | 2.91 | 0.65 | <.01 | 0.83 | |
| | Vv | Yes | 0.20 | 0.16 | 0.20 | 0.18 | |
| CG | | | | | | | 0.35 |
| | Dl | No | 0.34 | 0.10 | <.01 | 0.60 | |
| | Vv | No | -0.03 | 0.06 | 0.64 | -0.08 | |

*Note. N* = 10

**Supplementary Table 5.** *Spearman's Rho Between Behavioral Measures and Brain Optical Densities*

| Variable | *1* | *2* | *3* | *4* | *5* | *6* | *7* | *8* | *9* | *10* | *11* | *12* | *13* | *14* |
|---|---|---|---|---|---|---|---|---|---|---|---|---|---|---|
| 1. Distance | – | | | | | | | | | | | | | |
| 2. Velocity | 1.00* | – | | | | | | | | | | | | |
| 3. Top transitions | .87* | .87* | – | | | | | | | | | | | |
| 4. Time in top | .35 | .37* | .52* | – | | | | | | | | | | |
| 5. Latency to top | -.51* | -.52* | -.40* | -.42* | – | | | | | | | | | |
| 6. Time spent frozen | -.90* | -.89* | -.71* | -.36* | .60* | – | | | | | | | | |
| 7. Dm | .24 | .24 | .17* | .20 | -.07 | -.32 | – | | | | | | | |
| 8. Dl | .15 | .14 | .03 | .14 | .01 | -.23 | .87* | – | | | | | | |
| 9. Vs | .24 | .24 | .17 | .17 | -.22 | -.32 | .68 | .72* | – | | | | | |
| 10. Vv | .12 | .11 | .06 | -.03 | -.16 | -.18 | .41* | .33* | .37 | – | | | | |
| 11. POA | -.23 | -.23 | -.28 | .25 | -.15 | .16 | .31 | .29 | .48* | -.03 | – | | | |
| 12. Hb | -.03 | -.04 | -.14 | -.12 | .01 | -.19 | .42 | .42 | .47* | .43* | .09 | – | | |
| 13. Hc | -.11 | -.12 | -.04 | -.05 | -.03 | .04 | .39 | .40 | .35 | .43* | -.02 | .46* | – | |
| 14. CG | .02 | .00 | -.17 | -.11 | .05 | -.11 | .54 | .70 | .30 | .10 | .15 | .45* | .52* | – |

*Note. N*=63. **p* < .05

**Table 6. ROI measurements for each Brain Region**

| Brain Area | ROI Area (um) | Rostral to caudal range quantified |
|---|---|---|
| Dm | 5587.36 | 4 sections after appearance of central zone of ventral telencephalon until appearance of the habenula |
| Dl | 9877.66 | 4 sections after appearance of central zone of ventral telencephalon until appearance of the habenula |
| Vs | 3325.81 | Only at anterior commissure |
| Vv | 3238.51 | Starts at appearance of central zone of ventral telencephalon and ends at anterior commissure |
| POA | 3533.67 | Starts at the anterior commissure, stops 1 section before habenula |
| Hb | 1629.13 | Starts one section before disappearance of dorsal |

| | | |
|---|---|---|
| | | telencephalon until the periventricular gray zone of the optic tectem appear |
| Hc | 7685.74 | Starts when 5 sections after appearance of dorsal hypothalamus until pituitary disappears |
| CG | 1297.06 | Starts one section after end of central hypothalamus until disappearance of optic tectum |

**Table 7.** *Optical densities of each brain region in each fish.*

| ID | Strain | Treatment | Dm | Dl | Vs | Vv | POA | Hb | Hc | CG |
|---|---|---|---|---|---|---|---|---|---|---|
| 1 | Reactive | Stressed | 0.0008 | 0.0025 | 0.0207 | 0.0815 | 0.0587 | 0.0131 | 0.0037 | 0.0032 |
| 2 | Reactive | Stressed | 0.0170 | 0.0296 | 0.0467 | 0.0192 | 0.0951 | 0.0248 | 0.0793 | 0.0235 |
| 4 | Proactive | Stressed | 0.0432 | 0.0465 | 0.0590 | 0.0271 | 0.1176 | 0.0057 | 0.0085 | 0.0164 |
| 5 | Reactive | Stressed | 0.0015 | 0.0126 | 0.0197 | 0.0140 | 0.0261 | 0.0037 | 0.0172 | 0.0037 |
| 7 | Proactive | Stressed | 0.0067 | 0.0192 | 0.0670 | 0.0372 | 0.0214 | 0.0184 | 0.0369 | 0.0092 |
| 9 | Reactive | Stressed | 0.0075 | 0.0079 | 0.0249 | 0.0396 | 0.0654 | 0.0074 | 0.0621 | 0.0186 |
| 10 | Reactive | Stressed | 0.0027 | 0.0100 | 0.0118 | 0.0114 | 0.0561 | 0.0063 | 0.0191 | 0.0035 |
| 11 | Proactive | Stressed | 0.0402 | 0.0382 | 0.0504 | 0.0855 | 0.0236 | 0.0067 | 0.0786 | 0.0150 |
| 13 | Proactive | Stressed | 0.0038 | 0.0025 | 0.0265 | 0.0016 | 0.0322 | 0.0032 | 0.0221 | 0.0051 |
| 14 | Proactive | Stressed | 0.0169 | 0.0211 | 0.0513 | 0.0197 | 0.1194 | 0.0065 | 0.0149 | 0.0067 |
| 15 | Reactive | Stressed | 0.0006 | 0.0016 | 0.0386 | 0.0090 | 0.1885 | 0.0018 | 0.0128 | 0.0059 |
| 17 | Reactive | Stressed | 0.0043 | 0.0041 | 0.0074 | 0.0044 | 0.0247 | 0.0142 | 0.0271 | 0.0137 |

| | | | | | | | | | | |
|---|---|---|---|---|---|---|---|---|---|---|
| 19 | Proactive | Stressed | 0.0070 | 0.0065 | 0.0229 | 0.0035 | 0.0179 | 0.0048 | 0.0135 | 0.0053 |
| 21 | Proactive | Stressed | 0.0226 | 0.0191 | 0.0372 | 0.0843 | 0.0346 | 0.0219 | 0.0340 | 0.0061 |
| 22 | Proactive | Stressed | 0.0116 | 0.0089 | 0.0139 | 0.0053 | 0.0512 | 0.0071 | 0.0190 | 0.0180 |
| 23 | Proactive | Stressed | 0.0051 | 0.0034 | 0.0121 | 0.0028 | 0.0312 | 0.0091 | 0.0186 | 0.0105 |
| 24 | Reactive | Stressed | 0.0020 | 0.0029 | 0.0109 | 0.0020 | 0.0098 | 0.0018 | 0.0025 | 0.0101 |
| 25 | Reactive | Stressed | 0.0502 | 0.0412 | 0.0702 | 0.0341 | 0.1194 | 0.0266 | 0.0277 | 0.0175 |
| 26 | Proactive | Stressed | 0.0051 | 0.0067 | 0.0266 | 0.0010 | 0.0358 | 0.0072 | 0.0035 | 0.0085 |
| 27 | Proactive | Stressed | 0.0188 | 0.0193 | 0.0184 | 0.0209 | 0.0627 | 0.0040 | 0.0233 | 0.0063 |
| 28 | Proactive | Stressed | 0.0098 | 0.0059 | 0.0307 | 0.0130 | 0.0160 | 0.0034 | 0.0142 | 0.0059 |
| 29 | Proactive | Stressed | 0.0053 | 0.0121 | 0.0222 | 0.0162 | 0.0501 | 0.0095 | 0.0188 | 0.0090 |
| 30 | Reactive | Stressed | 0.0076 | 0.0122 | 0.0307 | 0.0177 | 0.0693 | 0.0022 | 0.0142 | 0.0059 |
| 33 | Proactive | Baseline | 0.0119 | 0.0171 | 0.1349 | 0.0207 | 0.0086 | 0.0075 | 0.0174 | 0.0067 |
| 34 | Proactive | Baseline | 0.0073 | 0.0154 | 0.0605 | 0.0144 | 0.0444 | 0.0208 | 0.0230 | 0.0161 |
| 35 | Proactive | Baseline | 0.0038 | 0.0071 | 0.0177 | 0.0150 | 0.0076 | 0.0032 | 0.0190 | 0.0112 |
| 36 | Proactive | Baseline | 0.0135 | 0.0068 | 0.0095 | 0.0045 | 0.0164 | 0.0034 | -0.0107 | -0.0002 |
| 37 | Proactive | Baseline | 0.0027 | 0.0268 | 0.0252 | 0.0072 | 0.0097 | 0.0152 | 0.0130 | 0.0152 |
| 38 | Proactive | Baseline | 0.0265 | 0.0031 | 0.0305 | 0.0078 | 0.0958 | 0.0079 | 0.0205 | 0.0081 |
| 39 | Proactive | Baseline | 0.0060 | 0.0257 | 0.0626 | 0.0481 | 0.0103 | 0.0461 | 0.0143 | 0.0067 |
| 40 | Proactive | Baseline | 0.0166 | 0.0086 | 0.0291 | 0.0055 | 0.0654 | 0.0074 | 0.0082 | 0.0065 |
| 41 | Proactive | Baseline | 0.0229 | 0.0177 | 0.0285 | 0.0094 | 0.0098 | 0.0028 | 0.0116 | 0.0038 |
| 42 | Proactive | Baseline | -0.0033 | 0.0363 | 0.0677 | 0.0411 | 0.0618 | 0.0150 | 0.0237 | 0.0105 |
| 43 | Proactive | Baseline | 0.0058 | 0.0086 | 0.0030 | -0.0063 | -0.0047 | -0.0017 | 0.0035 | -0.0002 |
| 44 | Proactive | Baseline | 0.0102 | 0.0102 | 0.0184 | -0.0027 | -0.0064 | 0.0005 | 0.0426 | 0.0069 |
| 45 | Proactive | Baseline | 0.0134 | 0.0179 | 0.0248 | 0.0062 | 0.0330 | 0.0138 | 0.0235 | 0.0213 |
| 46 | Proactive | Baseline | 0.0147 | 0.0200 | 0.0220 | 0.0047 | 0.0358 | 0.0158 | 0.0227 | 0.0212 |
| 47 | Proactive | Baseline | 0.0010 | 0.0326 | 0.0629 | 0.0111 | 0.0393 | 0.0014 | 0.0068 | 0.0083 |
| 48 | Proactive | Baseline | 0.0054 | 0.0020 | 0.0184 | 0.0094 | 0.0284 | 0.0035 | 0.0145 | 0.0105 |
| 49 | Reactive | Baseline | 0.0034 | 0.0049 | 0.0783 | 0.0409 | 0.0020 | 0.0087 | 0.0672 | 0.0393 |
| 50 | Reactive | Baseline | 0.0014 | 0.0024 | 0.0478 | 0.0086 | 0.0114 | 0.0101 | 0.0061 | 0.0119 |
| 51 | Reactive | Baseline | 0.0043 | 0.0015 | 0.0135 | 0.0068 | 0.0096 | 0.0041 | 0.0165 | 0.0050 |
| 52 | Reactive | Baseline | 0.0009 | 0.0045 | 0.0098 | 0.0146 | 0.0694 | 0.0074 | 0.0287 | 0.0160 |
| 53 | Reactive | Baseline | 0.0127 | 0.0005 | 0.0220 | 0.0009 | 0.0061 | 0.0078 | 0.0049 | 0.0157 |
| 54 | Reactive | Baseline | -0.0015 | 0.0169 | 0.0288 | 0.0133 | 0.0561 | 0.0219 | 0.0075 | 0.0057 |
| 55 | Reactive | Baseline | -0.0003 | -0.0007 | 0.0068 | 0.0024 | 0.0241 | -0.0022 | 0.0145 | 0.0118 |

| | | | | | | | | | | |
|---|---|---|---|---|---|---|---|---|---|---|
| 56 | Reactive | Baseline | 0.0053 | 0.0092 | 0.0060 | -0.0034 | 0.0180 | 0.0031 | 0.0092 | 0.0113 |
| 57 | Reactive | Baseline | 0.0176 | 0.0094 | 0.0707 | 0.0249 | 0.0218 | 0.0774 | 0.0166 | 0.0086 |
| 58 | Reactive | Baseline | 0.0089 | 0.0176 | 0.0871 | 0.0170 | 0.0374 | 0.0249 | 0.0050 | 0.0052 |
| 59 | Reactive | Baseline | 0.0040 | 0.0098 | 0.0132 | 0.0367 | 0.0653 | -0.0039 | 0.0188 | 0.0139 |
| 60 | Reactive | Baseline | 0.0213 | 0.0052 | 0.0082 | 0.0191 | 0.0436 | 0.0119 | 0.0350 | 0.0094 |
| 61 | Reactive | Baseline | 0.0129 | 0.0114 | 0.0477 | 0.0294 | 0.0342 | 0.0237 | 0.0171 | 0.0207 |
| 62 | Reactive | Baseline | 0.0091 | 0.0197 | 0.0382 | 0.0355 | 0.0966 | 0.0108 | 0.0403 | 0.0181 |
| 63 | Reactive | Baseline | 0.0059 | 0.0093 | 0.0195 | 0.0146 | 0.0314 | 0.0157 | 0.0035 | 0.0042 |

*Note. N = 63.*

**Supplementary Table 8**. *Quantification of discrete behaviors and composite score.*

| Stress Coping Style | Distance (cm) | Velocity (cm/s) | Top Transitions | Time in Top (s) | Latency to Top (s) | Not Moving (s) | PC1 Score | ID |
|---|---|---|---|---|---|---|---|---|
| Proactive | 9426.98 | 5.25865 | 130 | 762.095 | 0 | 0 | | 25 |
| Proactive | 9752.63 | 5.48657 | 119 | 629.096 | 2.1021 | 0 | | 28 |
| Proactive | 9004.23 | 5.00514 | 152 | 552.486 | 0 | 4.87154 | | 24 |
| Proactive | 10614.1 | 5.8974 | 296 | 981.582 | 42.309 | 0 | | 20 |
| Proactive | 10152.8 | 5.72731 | 249 | 893.393 | 16.5832 | 0 | | 15 |
| Proactive | 9422.75 | 5.23768 | 123 | 702.536 | 0 | 15.7491 | | 17 |
| Proactive | 11132.4 | 6.22889 | 251 | 515.516 | 8.07474 | 0 | | 12 |
| Proactive | 13131.1 | 7.3128 | 310 | 652.619 | 0 | 0 | | 13 |
| Proactive | 6038.54 | 3.38885 | 102 | 1327.93 | 86.1195 | 70.2035 | | 11 |
| Proactive | 8995.2 | 5.0264 | 189 | 859.293 | 0 | 0 | | 10 |
| Proactive | 9452.46 | 5.26787 | 132 | 677.911 | 0 | 73.6069 | | 9 |
| Proactive | 5516.6 | 3.07144 | 90 | 507.808 | 32.0654 | 214.982 | | 7 |
| Proactive | 9134.99 | 5.0882 | 211 | 980.247 | 0 | 0 | | 1 |
| Proactive | 9286.46 | 5.16249 | 174 | 797.164 | 0.033366 | 0 | | 2 |
| Proactive | 10165.5 | 5.73153 | 215 | 618.752 | 3.33667 | 0 | | 3 |
| Proactive | 5648.98 | 3.23017 | 151 | 1027.83 | 2.26893 | 0.633968 | | 4 |
| Reactive | 6718.61 | 3.73824 | 41 | 87.9213 | 0 | 134.001 | | 29 |
| Reactive | 4954.18 | 2.75513 | 42 | 82.1154 | 10.4104 | 234.268 | | 30 |
| Reactive | 3361.06 | 1.95906 | 15 | 24.5913 | 148.315 | 972.259 | | 27 |

| | | | | | | | | |
|---|---|---|---|---|---|---|---|---|
| Reactive | 2080.54 | 1.3186 | 48 | 209.71 | 2.66934 | 1313.13 | | 26 |
| Reactive | 3800.95 | 2.1246 | 123 | 533.4 | 503.47 | 928.128 | | 21 |
| Reactive | 2065.86 | 1.15011 | 6 | 1.66833 | 9.07574 | 1388.09 | | 23 |
| Reactive | 1566.03 | 1.33658 | 49 | 1385.084 | 4.63797 | 1433.032 | | 22 |
| Reactive | 3116.92 | 1.7349 | 19 | 154.988 | 1.1011 | 955.589 | | 19 |
| Reactive | 3058.61 | 1.84206 | 76 | 1347.05 | 21.7217 | 1203.5 | | 16 |
| Reactive | 1520.63 | 0.851424 | 4 | 2.03537 | 1108.78 | 1515.18 | | 14 |
| Reactive | 1648.77 | 0.916085 | 9 | 43.1431 | 989.923 | 1583.22 | | 18 |
| Reactive | 8527.88 | 4.82419 | 92 | 101.635 | 96.8302 | 0 | | 6 |
| Reactive | 5748.02 | 3.22117 | 96 | 516.483 | 10.01 | 196.096 | | 8 |
| Reactive | 1219.71 | 0.677678 | 25 | 61.6283 | 482.216 | 1633.1 | | 5 |

*Note. N = 30.*

**Supplementary Materials and Methods**

**Tissue Processing**: Brains were cryosectioned at 16 µm thickness onto four serial series. All slides were simultaneously post-fixed in cold 4% paraformaldahyde/PBS solution, washed in PBS and acetylated in 0.25% acetic anhydride/triethanolamine. Subsequently, slides were washed in 2X standard saline citrate, dehydrated in increasing ethanol series and stored at -80 C. One series was used to localize and quantify *egr-1* expression using a digoxigenin (DIG) labeled probe (see below).

**Probe Synthesis**: Digoxigenin (DIG) labeled *egr-1* riboprobes were generated using a 1:3 ratio of UTP and DIG-UTP (Roche) following a modified manufacturer's protocol (Megascript T7/T3, Ambion). The 393 base pair *egr-1* probe template was subcloned into a pCR4-TOPO vector (Invitrogen) by using primer pair 5'-GCAAGTATCCTAACCGGCCA -3'and 5'-GGTAGCTGGACACTGGTGAG -3'on the *D. rerio egr-1* transcript (Gene Accession NM_131248.1). After probe synthesis, we removed unincorporated nucleotides via column filtration according to manufacturer's protocol (Megaclear, Ambion) and checked probe quality through gel electrophoresis and quantity using a Nanodrop One (ThermoFisher).

***in situ* hybridization**: We used a modified digoxigenin *in situ* hybridization protocol [48,49]. Briefly, slides were prehybridized with solution containing 50% formamide, 5X SSC, 5X Denhardt's solution, 250 ug/ml yeast tRNA, and 500 ug/ml herring sperm DNA for 5 hours at 60°C in a hybridization chamber containing chamber buffer solution (50% formamide, 2X SSC). We then hybridized the slides 16 hours at 60°C with fresh prehybridization solution containing 600 ng of antisense riboprobe per slide in hybridization chambers containing buffer. Following hybridization we washed slides (2X SSC at room temperature, followed by 50% formamide in 2X SSC at 60°C, and then 2X SSC at room temperature), then RNase A treated the slides (0.5M NaCl, 10 mM Tris pH 8.0, 2.25 mM EDTA, 0.2 µg/ml RNase A), followed by increasingly stringent washes (2X, 1X, 0.5X, 0.25X SSC) and then a final wash in Buffer B1 (100 mM Tris pH 7.5, 150 mM NaCl). Sections were incubated 22 hours at 4°C with Anti-Digoxigenin AP antibody (Roche). After antibody incubation we washed sections twice in Buffer B1 and then blocked endogenous alkaline phosphatase activity with a 30 minutes wash in the dark in Buffer B3 (100mM Tris pH 9.5, 100 mM NaCl, 50 mM $MgCl_2$, 5 mM levamisole). We carried out colorimetric detection using NBT/BCIP stock solution (Roche). We stopped the colorimetric reaction after 11.3 hours by rinsing sections three times in ultrapure type 1 water, then progressively dehydrated sections in ethanol washes (25%, 50%, 70%, 95%), and finally coverslipped the slides with Permount adhesive (Fisher). All individuals were processed simultaneously to avoid any potential colorimetric development differences across individuals due to batch effects. DIG labeled *egr-1* sense riboprobes showed no expression.
